# Uniform processing and analysis of IGVF massively parallel reporter assay data with MPRAsnakeflow

**DOI:** 10.1101/2025.09.25.678548

**Authors:** Jonathan D. Rosen, Arjun Devadas Vasanthakumari, Kilian Salomon, Nikola de Lange, Pyaree Mohan Dash, Pia Keukeleire, Ali Hassan, Alejandro Barrera, Martin Kircher, Michael I. Love, Max Schubach

**Affiliations:** Department of Genetics, University of North Carolina, Chapel Hill, NC, USA; Institute of Human Genetics, University Hospital Schleswig-Holstein, University of Lübeck, Lübeck, Germany; Berlin Institute of Health at Charité – Universitätsmedizin Berlin, Berlin, Germany; Department of Biostatistics and Bioinformatics, Duke University Medical School, Durham, NC, USA; Center for Advanced Genomic Technologies, Duke University, Durham, NC, USA; Department of Biostatistics, University of North Carolina, Chapel Hill, NC, USA

## Abstract

As researchers and clinicians seek to identify human genomic alterations relevant to traits and disorders, identifying and aggregating evidence providing mechanistic support for associations between alterations and phenotypes remains challenging. In particular, the study of non-coding genomic variation remains a major challenge due to the lack of accurate functional annotation for activity in a given context and across alleles. Experimental evidence is critical for prioritizing and interpreting functional effects of genetic alterations. Massively Parallel Reporter Assays (MPRAs) have emerged as a powerful high-throughput approach, enabling quantification of regulatory element activity and allelic effects, and systematic dissection of gene regulatory logic and variant effects across different contexts. However, the diversity of MPRA designs, lack of standardized formats, and many potential processing parameters hamper data integration, reproducibility, and meta-analyses across studies.

To address these challenges, the Impact of Genomic Variation on Function (IGVF) Consortium established an MPRA focus group to develop community standards, including harmonized file formats, and robust analysis pipelines for a wide range of library types and experimental designs. Here, we present these formats and comprehensive computational tools, MPRAlib and MPRAsnakeflow, for uniform processing from raw sequencing reads to counts, processing and visualization. Using diverse MPRA datasets, we characterize technical variability sources including barcode sequence bias, outlier barcodes, and delivery method (episomal vs. lentiviral). Our results establish best practices for MPRA data generation and analysis, facilitating robust, reproducible research and large-scale integration. The presented tools and standards are publicly available, providing a foundation for future collaborative efforts in regulatory genomics.

## Introduction

Control of gene expression is a complex, dynamic process in which many regulatory elements determine when, where, and to what extent genes are active. Shifts in these regulatory programs drive development and shape evolutionary diversity, whereas disruptions can lead to disorders such as cancer, congenital malformations, and bleeding or metabolic disorders (Huang et al. 2013; VanderMeer and Ahituv 2011; Reijnen et al. 1992, 1993; Ludlow et al. 1996; De Castro-Orós et al. 2011). Cis-regulatory elements (CREs) such as enhancers and promoters modulate transcription through their recruitment of transcription factors and the surrounding chromatin landscape. Candidate CREs (cCREs) can be difficult to recognize and interpret because they may lack clear sequence or epigenetic hallmarks (e.g. open chromatin state or adjacent histone modifications), can influence genes over long genomic distances, and can have cell-type-specific activity. Consequently, pinpointing functional variants that drive phenotypic differences within and between species remains a major challenge.

The National Human Genome Research Institute (NHGRI) launched the Impact of Genomic Variation on Function (IGVF) Consortium in 2021, aiming to systematically understand how genomic variation affects genome function and how these effects shape phenotypes (Engreitz et al. 2024). Massively Parallel Reporter Assays (MPRAs) are a specific type of Multiplex Assays of Variant Effects (MAVEs) that can probe non-coding sequence effects and are a central approach within the IGVF efforts. MPRAs are high-throughput experiments that enable the simultaneous assessment of thousands to millions of genetic elements or variants in a single experiment. By inserting elements or variants into reporter constructs, one can measure their effects on diverse cellular processes like transcriptional initiation, splicing, transcript stability or protein function. To measure the effects of a cCRE on transcriptional activation, the test element is placed upstream of a minimal promoter and barcodes uniquely mapped to that test element allowing regulatory activity to be quantified by measuring the RNA levels of the transcribed barcode compared to the abundance of the barcode in DNA. Reporter plasmids can be tested episomally (episomalMPRA) (Melnikov et al. 2012), or integrated into the genome using lentiviral delivery (lentiMPRA) (Inoue et al. 2017), transposons or recombination. While these experiments enable testing thousands of elements in parallel, both episomal and genomic integrated MPRAs measure regulatory activity outside of the native genomic locus and thus may not fully recapitulate *in vivo* regulatory dynamics (Kosicki et al. 2025).

A related method, Self-Transcribing Active Regulatory Region sequencing (STARR-seq), can be used simultaneously to study both cCREs and variants (Arnold et al. 2013; van Arensbergen et al. 2017). Unlike standard MPRAs, STARR-seq places cCREs downstream of a minimal promoter, allowing each element to serve as its own transcribed barcode and permitting direct quantification of enhancer activity based on the abundance of self-transcribed RNA. While STARR-seq assays are also conducted within the IGVF consortium, differences in assay outputs and data processing necessitate different processing. In this work, the primary focus is on barcode-based MPRAs, with STARR-seq included or referenced where relevant.

In recent years, there has been a substantial increase in MPRA publications, with laboratories worldwide employing the technique to characterize cCREs or identify expression altering variants (Kircher et al. 2019; Jiang et al. 2024; Agarwal et al. 2025). MPRAs are being used to study variants in genomic cohorts (Koesterich et al. 2023), to develop predictive tools (Agarwal et al. 2025; de Almeida et al. 2022; Gosai et al. 2024), as benchmarks (Jaganathan et al. 2025; Avsec et al. 2021; Linder et al. 2025; Avsec et al. 2025; Rafi et al. 2024), and to validate synthetic sequences designed for specific behaviors such as tissue-specific expression. These applications illustrate that MPRAs are not only interpreted in isolation, but now form the basis for numerous meta-analyses. Accordingly, standards for MPRA experiments are essential to enable the re-analysis and integration of data from diverse studies as well as access for a growing community. To address this need, the IGVF consortium established an MPRA focus group in 2021 to develop community standards, file formats, and pipelines for consistent data processing and experimental coordination across the consortium (Engreitz et al. 2024).

Here, we present key products of this focus group, enabling researchers to conduct and analyze MPRA studies using standardized approaches. We describe critical aspects of MPRA design, introduce standardized file formats for reporting both assay design and results, and present our newly developed MPRA processing pipeline MPRAsnakeflow. This pipeline supports uniform analysis of diverse MPRA assays from sequencing reads to count data. Notably, we optimized MPRAsnakeflow for the increasing number of variant designs and demonstrate that previously used strategies can perform poorly due to issues arising from small edit distances in designed sequences. Complementing MPRAsnakeflow, we developed a Python library, MPRAlib, which facilitates downstream processing of DNA and RNA counts, filtering, outlier removal, and data visualization, giving researchers a powerful tool for integrating MPRA data into python notebooks and linking to other statistical tools (Ashuach et al. 2019; Keukeleire et al. 2025; Myint et al. 2019). We analyze multiple assays from IGVF and ENCODE, comparing barcode discrepancies and outliers across replicates and experiments. We discuss GC effects, outlier consistency across replicates, and differences between episomal and lenti-integration based MPRAs. We note that assays performed in heterogeneous cell cultures, such as those undergoing differentiation, exhibit a greater number of barcode outliers. In summary, we provide a comprehensive set of tools for MPRA data processing and highlight important considerations for robust and reproducible MPRA analysis.

## Methods

### MPRA design

Arbitrary sequences can in principle be tested via an MPRA, constrained only by technical limitations like construct or vector length. Typically, sequences are generated by DNA oligo synthesis, which currently is still most economical for sequence lengths around 300 nucleotides (nts) and covers a majority of open chromatin peak lengths. Longer sequences can either be derived from combining shorter sequences or from selections or amplifications of genomic DNA fragments. When barcodes are not directly synthesized with each sequence but added randomly (e.g. by amplification with overhanging barcode primers), the resulting MPRA library must be sequenced to determine barcode-to-test sequence associations. Paired-end short read sequencing is usually sufficient to cover the entire test sequence with a small overlap (we recommend >10 nts for high-complexity sequences). For example, 2×150 nt paired-end sequencing can cover a 200 nt test oligo + 20 nt barcode + 30 nt spacer with 50 nt overlap.

Alternatively, long-read sequencing can be employed. However, we note that various MPRA designs have different requirements on the precision of these sequence read-outs to obtain an unambiguous association, specifically if sequence variants need to be distinguished. Some MPRA experiments test synthetic or engineered sequences that possess particular activity properties, such as tissue-specific enhancer activity or optimal spacing between two elements (de Almeida et al. 2022; Gosai et al. 2024; Georgakopoulos-Soares et al. 2023). However, most MPRAs are used to test candidate cis-regulatory elements (cCREs) derived from a genomic or inferred (ancestral) genomic sequence. The selection process generally follows two complementary strategies:

Phenotype-driven design – In a phenotype-driven approach, variants or regulatory elements are selected based on known disease-associated genes or pathways (Castaldi et al. 2019). This design emphasizes proximity to implicated genes or loci, or association via other biochemical experiments. Candidate elements may include the gene’s promoter, nearby distal enhancer elements suggested by chromatin maps, or regions containing variants associated with disease from genome-wide association studies (GWAS) or other prior studies (Deng et al. 2024; McAfee et al. 2023). Sequences with known transcription factor binding sites (TFBSs) or other biochemical marks (e.g., open chromatin identified by DNase- or ATAC-seq peaks) may also be prioritized to increase the likelihood of regulatory activity (Agarwal et al. 2025).

Genotype-driven design – In a genotype-driven design, individual genetic variants are the starting point. Variants of interest include GWAS tag single nucleotide polymorphisms (SNPs), high-confidence and fine-mapped candidate expression quantitative trait loci (eQTLs), or rare and singleton variants identified in disease cohorts or genome projects (e.g., de novo variants in ASD (Koesterich et al. 2023), gnomAD database (Chen et al. 2024)). For each variant, oligonucleotides are designed to span the variant site (Farrow et al. 2024; Abell et al. 2022). If multiple variants are in linkage disequilibrium (LD) within the same region, separate oligos for each haplotype can be synthesized or non-target positions can be fixed to the major allele (Abell et al. 2022).

For both strategies, cCREs are typically identified using publicly available genomic annotations. Proximity to genes and regulatory elements (promoters or enhancers) can be used to prioritize cCREs for testing. High-resolution chromatin contact maps (Hi-C, capture Hi-C) can further pinpoint physical interactions with promoters or distal enhancers (Thakur et al. 2024). Candidate sequences may also be selected for TFBS, i.e. sequences containing motifs (known or predicted) or overlapping ChIP-seq peaks for relevant transcription factors to enrich for regulatory activity. At the promoter level, candidate sequences may include transcription start sites (TSSs) and proximal upstream motifs (e.g., *de novo* promoter libraries). Sometimes tiling of longer sequences might be required to capture variable regulatory grammar (Kosicki et al. 2025).

In addition to test sequences (e.g., variants or cCREs), control sequences are required to validate experiment performance. By allowing the quantification of baseline activity, they help to identify significantly activating or repressing sequences and establish that the activity distribution of tested sequences exceeds that of negative controls. The most common approach is to use shuffled sequences as negative controls to estimate background activity. Positive element or variant controls are more challenging to design. Ideally there exist previously tested sequences with high activity in the same cell type, or variants with known properties in the same system. If such sequences are not available, sequences with activity across many cell-types or from a related cell-type should be used. Generally, only a small number of positive controls (around hundred) might be required while a larger set of several hundred negative controls bolsters later statistical testing. However, if previously validated positive controls are unavailable, more putative positive controls are recommended.

We recommend avoiding the use of sequences with certain undesirable properties. Sequences containing long homopolymers or specific restriction sites (e.g., EcoRI or SbfI) used in MPRA library construction should be removed. Restriction sites may also be eliminated by sequence modification. Simple repeats and overlap with transposable elements may cause unwanted strong activity or construct silencing. The inclusion of sequences with binding sites for factors impacting chromatin structure (e.g. CTCF binding sites) should be excluded, since short MPRA constructs cannot model chromatin looping structures. Finally, sequences containing additional TSSs may interfere with the minimal promoter, resulting in inconsistent background activity of the core element.

To facilitate this process, we developed a workflow called MPRAOligoDesign (https://github.com/kircherlab/MPRAOligoDesign), which can generate a design file from sequences, variants, genomic regions, and paired variants with genomic regions. It extends regions to the desired sequence length, tiles longer regions, designs reference and alternative sequences for variants, and can be configured to perform recommended property checks, removing sequences that do not pass the filters.

### Standardized file formats

Within the last ten years, over 100 MPRA experiments have been published. Various efforts have been made to integrate different experiments into common databases, like MaveDB, MPRAVarDB, and MPRAbase (Rubin et al. 2025; Jin et al. 2024; Zhao et al. 2025). However, since there are no defined standards for metadata, such as the genomic region tested or sequence design, setting up similar data resources becomes very challenging. One major resource of IGVF will be a catalog of element and variant effects. With various different MPRAs being conducted, they must be integrated into a single catalog as well as a data portal for reuse by other projects. We see a tremendous need to harmonize metadata regarding the design and various stages of output files for MPRA data. We identified four categories for which standardized files are necessary: Design, Counts, Effects, and Genomic effects, and provide a detailed documentation of the file formats in Supplementary Note S1.

For “Design”, we defined a reporter sequence design file that contains the main metadata of designed oligos such as identifier, sequence, sequence origin and class and provides key information about the purpose of the assay. Despite the “designed oligo” terminology, such design files should also be created for MPRA experiments not derived from oligo synthesis and equivalent information should be collected. In the design file, each oligo has a unique name or ID and the nucleotide sequence. It further includes the category (variant, element, synthetic, scrambled), an assigned class (test, variant positive control, variant negative control, element active control, element inactive control), and additional information. Chromosome, start and end positions, along with the reference genome version, must be specified if applicable. For variants, the position within the tested sequence, a unique ID using the canonical SPDI notation (Holmes et al. 2020), and whether the reference or alternative allele was tested within the sequence must also be included. This file enables re-analysis of the data as well as the generation of element and/or variant effects from the data.

“Count” files report the number of observed reads (i.e. integer counts) per barcode or aggregated to elements per modality (i.e. DNA or RNA). Multiple counts and modalities can be described in the same file. Barcode-level files contain the barcode sequence and the associated oligo identifier. Element-aggregated files describe the replicate ID, oligo ID, aggregated and normalized DNA and RNA counts, the number of aggregated barcodes and an inferred log-fold change (normalized RNA/DNA ratio). For IGVF, these are generated via the standardized pipeline MPRAsnakeflow (see below). However, existing published pipelines, like MPRAflow (Gordon et al. 2020), also generate similar types of data and could be converted to our standardized format.

“Effect” files report the results of differential activity analysis. This includes normalized counts, activity estimates (i.e. log-fold changes) and significance estimates (i.e. p- and q-values). This can be performed using tools like MPRAnalyze, BCalm, or mpralm (Ashuach et al. 2019; Keukeleire et al. 2025; Myint et al. 2019), with the count data and information from the sequence design file as input (e.g., to map oligos to reference or alternative alleles for a variant). Because differential activity analysis differs between elements and variants (for example, statistical testing for elements always requires a control set of oligos), we separate them into element and variant effects files. In addition to the values listed above, element files contain aggregated modality counts. Variant files contain those for both the alternative and reference allele, a posterior probability of the regulatory effect, and a 95%-confidence interval of the effect.

Finally, “Genomic effects” are those that can be mapped to a genomic location, and we have created standardized BED files for variants and elements. These contain information from effect files with genomic coordinates for visualization of MPRA results in genome browsers such as UCSC or IGV (Perez et al. 2025; Robinson et al. 2011). Tools for MPRA data from IGVF will support these standards.

In our MPRAlib library (see below) a command set called `validate-file` is implemented to compare actual files with their schemas to ensure files are formatted correctly.

### A unified MPRA pipeline: MPRAsnakeflow

MPRAsnakeflow is a further evolution of MPRAflow (Gordon et al. 2020), which established basic functionality for the analysis of MPRA data. We focus on six topics to enhance and extend its functionality: (1) Speed improvements to support large complex MPRA assays, (2) Interoperability by containerization of software dependencies, (3) Standardized outputs, quality reports and additional statistics to help uncover potential issues with the MPRA experiment, (4) Support for multiple assays (incl. raw read structures), (5) Optimized barcode assignment to support designs with small edit distances, like multiple variants for the same cCRE, and (6) New features like strand awareness, outlier removal and support of multiple tools for statistical quantification. MPRsnakeflow is implemented in the workflow language Snakemake (Mölder et al. 2021). It is organized into two sub-workflows: the assignment workflow and the experiment workflow (see Figure 1).

**Figure 1:**
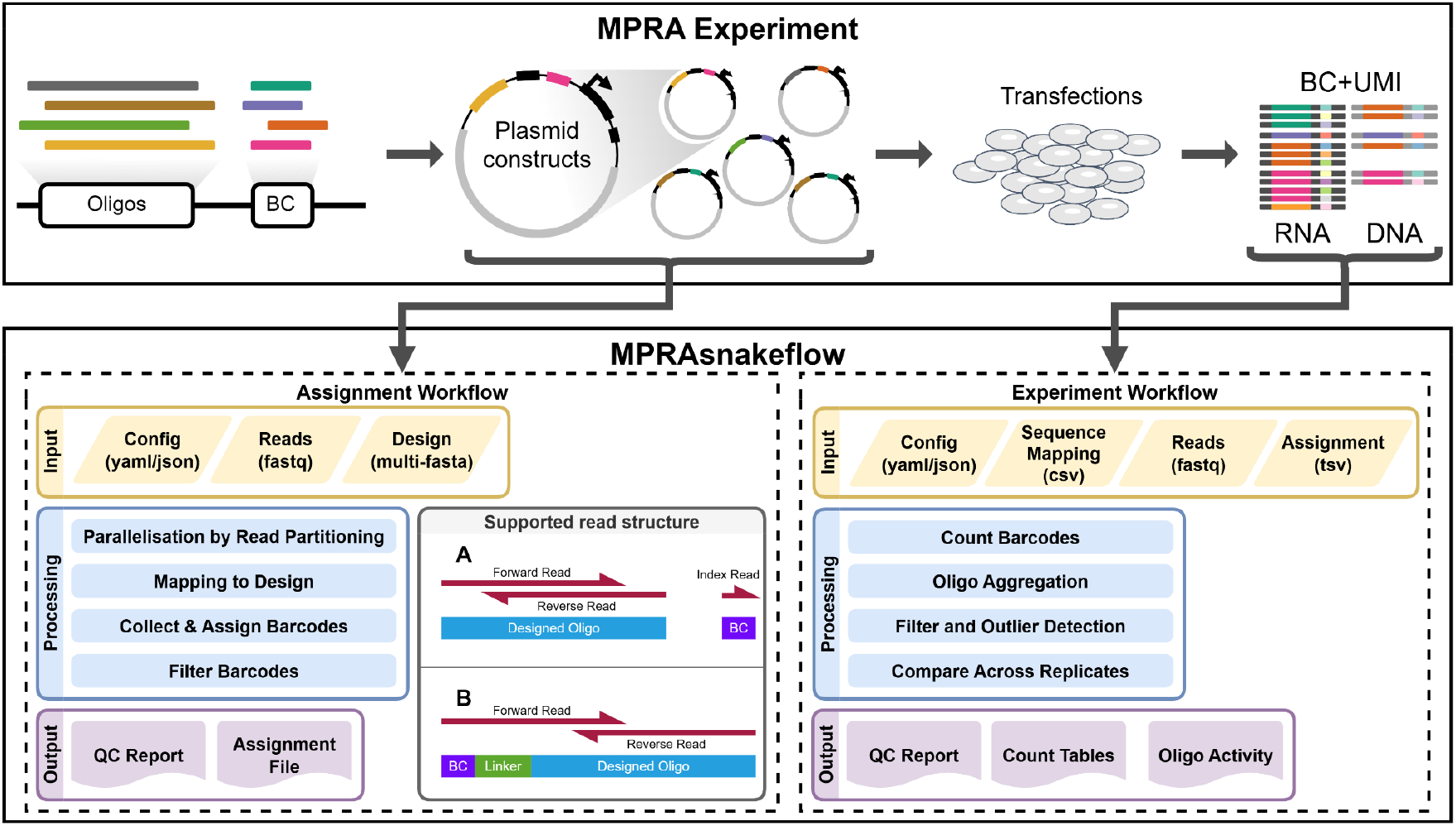
Overview of an MPRA experiment with UMIs (top) and the MPRAsnakeflow pipeline for analyzing MPRA experiments (bottom). MPRAsnakeflow consists of an assignment workflow (left), which maps barcodes to designed oligos, and an experiment workflow (right), which counts barcodes and computes oligo activity. Inputs are shown in yellow, main processing steps in blue, and outputs in purple. Notably, MPRAsnakeflow supports different input read structures, including separate reads for oligo and barcode (A), as well as combined structures where barcode and oligo are present within read pairs (B).

The assignment workflow aims to associate barcodes with designed oligos. It supports multiple read structures typical for certain assay types (Gordon et al. 2020; Gosai et al. 2024). For example, the barcode can either be sequenced in a separate index read or positioned at the beginning of the first paired-end read, separated from the start of the oligo by a linker. In brief, the assignment workflow first verifies that the design file (in multi-FASTA format) is appropriate, ensuring unique headers and no duplicate sequences. Next, paired-end reads of the sequenced oligos are merged using NGmerge (Gaspar 2018), with a minimum recommended overlap of 11 nts. The merged reads are then mapped to the design file using a read mapper (BBmap (Bushnell 2014) [default] or BWA-MEM (Li and Durbin 2009)) or by exact string matching. Barcodes are assigned based on the best unique sequence match. Finally, barcodes are aggregated and filtered based on a minimum number of required observations per oligo (default: 3), and a majority of barcode occurrences must map to the same oligo (default: 0.75) to minimize ambiguity while allowing barcode collisions or best alignment errors. This workflow can be skipped if barcode associations are already known (e.g., by design or using custom assignment scripts).

The experiment workflow processes the sequenced barcodes extracted from DNA and RNA samples of the MPRA experiment in a specific tissue or cell type. It counts the observations of each barcode, merges DNA and RNA counts, and assigns barcodes to oligos using an assignment file, which can be generated by the assignment workflow. An optional step removes outlier barcodes. Barcode counts per oligo are then aggregated, and activity is calculated as the log2 ratio of normalized (counts per million reads) RNA over DNA counts for each oligo. When multiple replicates are available, the workflow computes correlations of DNA counts, RNA counts, and log2 ratios to assess data quality. Additionally, the experiment workflow supports the use of UMIs to disambiguate unique transcripts from PCR duplicates to reduce PCR artifacts.

Both sub-workflows produce a variety of statistics and plots to help users inspect the data and identify potential errors. A unique feature of MPRAsnakeflow is the quality report generated for both workflows, which summarizes key metrics and plots along with additional explanations. The quality report is provided as a single structured and image-embedded HTML file, enabling quick and efficient data inspection and making it easy to share results with collaborators.

### Mapping strategies for barcode associations

To mitigate the impact of sequencing errors, sequenced oligos are typically mapped (Alser et al. 2021) to the originally designed sequences of the MPRA experiment. Approaches relying solely on exact string matching, as implemented in some other pipelines, like esMPRA (Li et al. 2025), fail to account for sequencing errors and consequently result in a reduced number of assigned reads and therefore less observed barcodes. Our observations indicate that the choice of mapping strategy can substantially influence downstream analyses, depending on the library design. For this reason, MPRAsnakeflow offers multiple strategies: exact matching (requiring the full read sequence matching the designed oligo, in either the forward or reverse-complement orientation), BWA-MEM or BBMap.

We utilize the BWA maximal exact matches (MEM) algorithm (Li and Durbin 2009) with an increased clipping penalty (default: 80) to discourage soft-clipping of read-ends by the aligner, effectively requiring a global sequence alignment. The mapper calculates a mapping quality score (MAPQ) based on the difference between the best and the next-best alignment. In MPRAsnakeflow, BWA-MEM alignments are then filtered using a configurable MAPQ threshold (default: 1) to remove ambiguous mappings. We observed that BWA-MEM reports MAPQ 0 values despite distinguishing oligos with minimal sequence differences in alignment (as apparent from alignment score differences reported in the output). This causes barcodes to be lost from associations in variant-rich MPRA libraries. To address this, we implemented an additional filtering step for BWA alignments, allowing users to still consider MAPQ 0 results based on sequence similarity, number of mismatches, and alignment length.

Additionally, MPRAsnakeflow provides support for BBMap (Bushnell 2014), an alternative k-mer-based aligner. BBMap reports higher MAPQ values, reflecting increased alignment confidence even for variant-rich libraries. For BBMap, a more stringent default MAPQ threshold (30) is applied to remove ambiguous assignments.

To compare the performance of these mapping strategies, we analyzed three distinct MPRA datasets of different library sizes and design choices: 8K-neurons, 20K-HepG2_P, and 80K-neurons (see Supplementary Note S2 and Supplementary Table S1 for a description of the datasets). For each dataset, we applied the different mapping routines and compared the number of assignable oligos, the identified genomic regions, and the number of variant sequences detected (see Figure 2). Only oligos associated with at least 10 unique barcodes were considered for these analyses. Furthermore, we used BCalm (Keukeleire et al. 2025) to obtain a variant effect quantification (log2 fold change) and a Benjamini-Hochberg adjusted p-value per variant, omitting outlier barcodes, to identify significant variant effects (adj. p-value < 0.1), and to assess the consistency of results across the tested mapping strategies.

**Figure 2:**
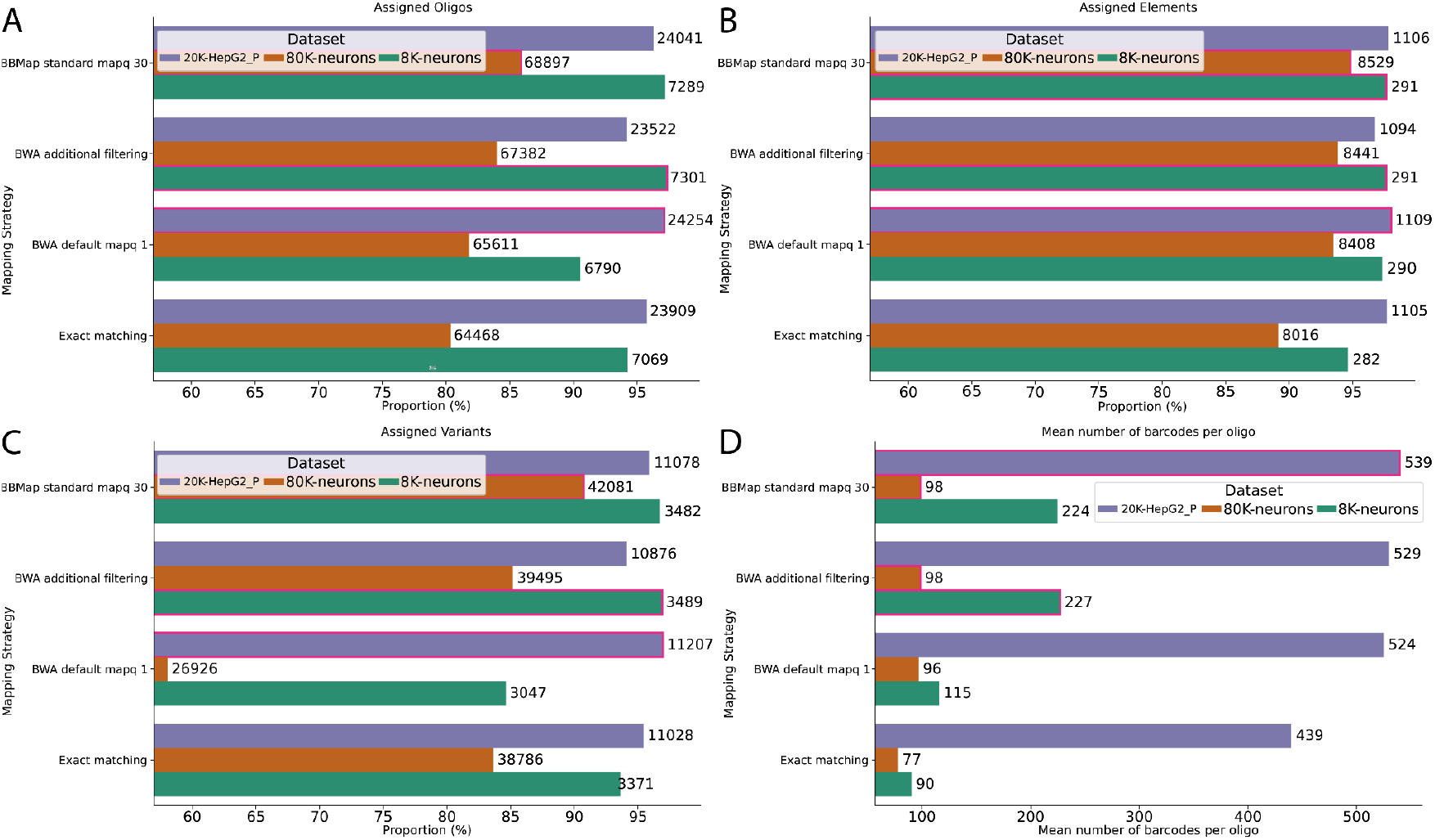
Comparison of four different mapping strategies for the association step for three different MPRA libraries (20K-HepG2_P, 80K-neurons, 8K-neurons). (A) Association of oligos, (B) Association collapsed to elements (cCREs), or (C) Association of variants. A variant is only considered if both the reference and the alternative allele oligos are assigned. (D) Mean number of barcodes per oligo. Bar with the best result for each MPRA library is highlighted in each panel.

### Strand sensitivity in the association step

Most read mappers consider alignments for both strands and therefore do not distinguish between the two possible orientations of a test sequence. However, MPRA constructs can have orientation-specific effects so libraries can contain oligos designed in both orientations. Depending on the preparation step for association sequencing, sequence orientation might be lost or maintained. Some MPRA designs test effects of sequence orientation and therefore maintain orientation information in the sequencing library. MPRAsnakeflow provides a strand-sensitive mode, which appends unique flanking sequences to both the reference design and the sequencing reads in the association step. This enables strand-specific analysis and allows for the systematic testing of promoters and enhancers in both orientations (Agarwal et al. 2025).

### Complexity and sequencing depth estimation

A common question is whether deeper sequencing of an MPRA library improves data quality, such as increasing correlation across replicates. To address this, the pipeline implements two key indicators. First, we use pairs of replicates to estimate overall library complexity using the Lincoln-Petersen method (Lincoln 1930; Petersen 1896). The difference between the observed number of barcodes and the Lincoln index provides an estimate of the potential improvement achievable with additional sequencing. The Lincoln index can also be used to identify replicates or modalities with decreased barcode complexity. Second, the pipeline includes a built-in downsampling option to assess whether quality metrics (e.g., replicate correlation) already reach saturation. If not, this indicates the potential benefit of further sequencing. Downsampling can be performed either on individual barcode counts or on barcodes within the assignment file, using either a fixed number or a specified proportion. Multiple downsampling configurations can be applied within a single experiment run using named configurations, ensuring that only the necessary parts of the workflow are rerun, which saves disk space and computational resources.

To demonstrate the utility of these approaches, we randomly downsampled DNA and RNA counts from 5% to 95% of the original counts, in 5% increments. This analysis was performed on libraries of varying complexity: low (8K-neurons), medium (80K-neurons), and high (240K-HepG2) complexity (see Supplementary Table S1). After downsampling, we computed the Lincoln index, the median number of barcodes per oligo, and the total number of oligos retained with at least 10 barcodes. To measure consistency of activity scores between replicates, we calculated Spearman correlation across replicates at each downsampling level. We compared these results to those obtained using the downsampling strategy implemented in MPRAlib (see below) to assess whether the metrics are comparable.

### A library for interactive MPRA count data analysis: MPRAlib

We developed a Python library, MPRAlib, to provide a simple and accessible framework for analyzing MPRA count data (Supplementary Figure S1). MPRAlib uses the AnnData object (Virshup et al. 2024) as its core data structure, with variables representing barcodes or oligos and observations corresponding to experimental replicates. The AnnData object is designed to support both barcode-level and oligo-level count data, enabling functionalities such as barcode outlier detection and data aggregation per oligo.

The main class, `MPRAData`, is an abstract base class that implements most core functions. Two subclasses, `MPRABarcodeData` and `MPRAoligoData`, inherit from `MPRAData` to support analyses at different data granularities. AnnData layers are used to store different modalities (e.g., RNA or DNA counts), normalized counts, and activity measures (such as log2 ratios). The flexible AnnData structure also allows the addition of metadata and pairwise data to variables or observations, which is leveraged to store information such as the number of barcodes per oligo, masking arrays for filtered data, and pairwise correlations across replicates.

Built-in functions are provided for reading MPRAsnakeflow outputs (at both barcode and oligo levels). We implemented a dynamic approach for adjusting the minimum required number of barcodes per oligo, allowing users to change this threshold at runtime without losing access to the original raw data. This flexibility enables users to recompute metrics, such as correlations, on different thresholds, without reloading the data. At the barcode level, MPRAlib offers various options for detecting outliers, setting minimum required counts per modality, and subsampling counts or barcodes. The library includes a comprehensive suite of plotting functions, such as replicate correlations, DNA versus RNA scatter plots, and histograms of barcodes per oligo.

To facilitate integration with differential analysis tools such as BCalm, mpralm, and MPRAnalyze (Keukeleire et al. 2025; Myint et al. 2019; Ashuach et al. 2019), MPRAlib supports a standardized metadata format and includes functions for exporting data in formats compatible with these tools. Additionally, MPRAlib can import results from these tools and generate further outputs, such as BED file tracks for genome browsers.

MPRAlib is extensively documented (https://mpralib.readthedocs.io) and is accompanied by several tutorials using Python notebooks, demonstrating its use in various scenarios. For convenience, we also provide a command-line interface (CLI) implementing commonly used workflows, such as correlation analyses and plotting routines. The CLI allows users to validate input file formats against standardized schemas, which are defined using JSON Schema (see Supplementary Note S1). The library is written in Python (version ≥3.9) and is available for installation via PyPI and Bioconda (Grüning et al. 2018).

### Barcode discrepancy detection and outlier removal

For the outlier analysis, seven IGVF and one ENCODE MPRA datasets were selected as follows. LentiMPRAs: 240K-HEK293T, 240K-HepG2, 12K-cardiop, 12K-cardiom, 80K-WTC11, 80K-neurons. EpisomalMPRAs: 20K-HepG2_P, 60K-A549_P. See Supplementary Note S2 and Supplementary Table S1 for more description on the datasets. Raw count data were processed to calculate counts per million (CPM) for both DNA and RNA libraries in each replicate. We computed the sum of CPM values across replicates for DNA (dna_cpm_sum) and RNA (rna_cpm_sum), followed by calculation of expression ratios as log2(rna_cpm_sum/dna_cpm_sum). Quality filtering was applied to remove low-confidence measurements: barcodes with zero RNA CPM or DNA CPM below the 5th percentile were excluded. Additionally, we retained only oligos represented by at least 20 barcodes across replicates to ensure sufficient statistical power. We implemented three complementary outlier detection approaches: **Global Outliers:** For each replicate, we calculated z-scores for RNA counts across all barcodes and flagged barcodes with absolute z-scores exceeding 3 as global outliers. **Oligo-Specific Outliers:** For each oligo, we computed the mean and standard deviation of RNA counts for each replicate. Barcodes deviating more than 3 standard deviations from their oligo-specific mean were classified as oligo outliers. Standard deviations of zero were replaced with 1 to handle oligos with invariant expression. **Large Expression Outliers:** We calculated the median expression ratio for each oligo and identified barcodes with expression ratios exceeding 5 log2-units above their oligo-specific median as large expression outliers. These outlier detection methods are also implemented in MPRAlib.

Outlier barcode sequences were analyzed for GC content, using the outlier sequences as the test sequences and non-outlier sequences identified from all replicates and all methods as a common background. Finally, the variant testing results of the MPRA experiments generated using BCalm (Keukeleire et al. 2025) were compared before and after removing outliers to check for any differences in both number and type of sequences that are identified as significant.

## Results

### Mapping strategies

Using different mapping strategies, we report the assignable oligos (Figure 2A), identified genomic regions (Figure 2B), the assigned variants (for which both reference and alternative oligo must be present, Figure 2C), as well as the number of barcodes per oligo (Figure 2D). In short, BBMap was able to assign more oligos compared to BWA-MEM and exact matching across all MPRA sets used. The same holds for assigned regions and especially for variants (Figure 2C). We see a drastic decrease in reported variants using the BWA mapping strategy, confirming the outlined issues with MapQ values where two oligos are in close edit distance (e.g., one nucleotide for SNVs). Apart from the different number of mapped oligos, we could not identify large effects on replicate correlation (n=3) across mapping strategies (see Supplementary Figure S2A and S2B).

We further inspected the design on 80K-neurons/WTC11 because of the difference in variants for BWA-MEM and noticed that most reference alleles had at least 3 alternative alleles each. Within this subset of oligos, the proportion of assignable variants dropped to below 40% (see Supplementary Figure S4). This shows that small edit distances play a crucial role in retaining variants and should be carefully considered for elements with many designed variants or for saturation mutagenesis (Kircher et al. 2019).

To compare the impact on downstream analyses, we utilized BCalm (Keukeleire et al. 2025) to determine significant variant effects for all four mapping strategies using two different MPRA experiments (8K-neurons, 80K-neurons). Supplementary Figure S3 shows the overlap in all variants and significant variants. BWA-MEM detected the lowest number of variants, followed by the exact matching, while most variants were either detected by the additional filtering applied to BWA output (8K-neurons, Supplementary Figure S3A) or BBMap (80K-neurons, Supplementary Figure S3B).

Variants detected by the default BWA-MEM were detected by all other mapping strategies, and only 0.1% (n=2, 8K-neurons) or 0.2% (n=96, 80K-neurons) of the variants were only in the exact matching strategy for the 8K-neurons and 80K-neurons dataset, respectively. Overall, the concordance in identified variants is very high, for example, the proportion of variants detected across at least three out of four methods is 97.6% (n=3406, 8K-neurons) and 94.7% (n=37956, 80K-neurons). When looking into the significant variants, concordance between the mapping strategies drops, with only 48.7% (n=416, 8K-neurons) and 59.2% (n=1011, 80K-neurons) of variants being significant for at least three out of four mapping strategies. Concordingly, we observe larger numbers of variants only significant for one of the mapping strategies. This occurs for 14.4% (n=123, 8K-neurons) of variants (Supplementary Figure S3C) and 24.3% (n=416, 80K-neurons) of variants (Supplementary Figure S3D), respectively. Notably for 8K-neurons, BWA-MEM with additional filtering (n=721) and BBMap (n=710) approaches yielded sets that are more than 1.5-fold larger than those of the other strategies (BWA n=456, exact matching n=420).

In summary, the choice of the mapping strategy for the association significantly influences variant analysis outcomes on the count level, particularly for complex and large libraries. Exact matching approaches fail to account for sequencing errors, leading to fewer barcode and oligo assignments, while error-tolerant methods like BWA struggle with confident distinction of related sequences, necessitating custom scripts for mitigating this issue. MPRAsnakeflow implements BBmap as standard, because it showed an overall good performance (Figure 2), but we recommend checking for other mapping strategies if the number of assigned barcodes are low.

### Complexity and sequencing depth estimation

For the small library experiment (8K-neurons, see Supplementary Table S1 for details about the design), we observe a median of 1,444,480 assigned barcodes across replicates after assignment and RNA and DNA merging, requiring barcodes to have at least one count of DNA and RNA in the full data set. The median of pairwise replicate Lincoln index values is 1,523,572. Consequently, we are missing around 5% of the barcodes from each replicate within this dataset. We assume that this might be a common loss from transfections or transductions of MPRA libraries into cell populations.

The medium MPRA library (80K-neurons, see Supplementary Table S1) presents a median of 5,459,247 assigned barcodes across replicates and a median Lincoln index of 6,243,618. Thus, we are missing approximately 12% of the library in each replicate. The Lincoln index for the large 240K-HepG2 library (see Supplementary Table S1) is 20,408,916 and we observe a median of 13,038,694 assigned barcodes across replicates. This is approximately 36% of missing barcodes.

Depending on the size of the designed library and the estimated complexity of the data, we see an increasing gap between missing barcodes in our libraries and a higher variability across replicates. For the medium and low complexity libraries, we see a good concordance between replicates, but the question arises whether more sequencing data could help for the higher complexity library. Therefore downsampling of DNA and RNA counts was performed for fixed proportions with MPRAsnakeflow (before assignment) and also of the final assigned count file (barcode reporter experiment) using MPRAlib. For all libraries (8K-neurons: Supplementary Figure S5A; 80K-neurons: Figure 3A and 3B; 240K-HepG2: Supplementary Figure S5B) we see a saturation in number of assigned barcodes as well as the observed oligos and the Pearson correlation of the oligo activity (log2 fold change). Both approaches show very similar trajectories and similar values for the observed number of barcodes. MPRAsnakeflow downsampling has a smoother curve and is more accurate but takes more time whereas using MPRAlib can be done within minutes, depending on the library size. Based on these results, 8K-neurons and 80K-neurons are already saturated and new sequencing data will not substantially improve these metrics (Figure 3 and Supplementary Figure S5A). For 240K-HepG2 (Supplementary Figure S5B) and, to a lesser extent, for 80K-neurons we still see a gently ascending line, but with no improvement in the number of detected oligos and Pearson correlation. Therefore the benefits of resequencing will be neglectable. Finally, we note that we are unable to retain all oligos with at least 10 barcodes because the underlying assignment contains oligos with fewer barcodes.

**Figure 3:**
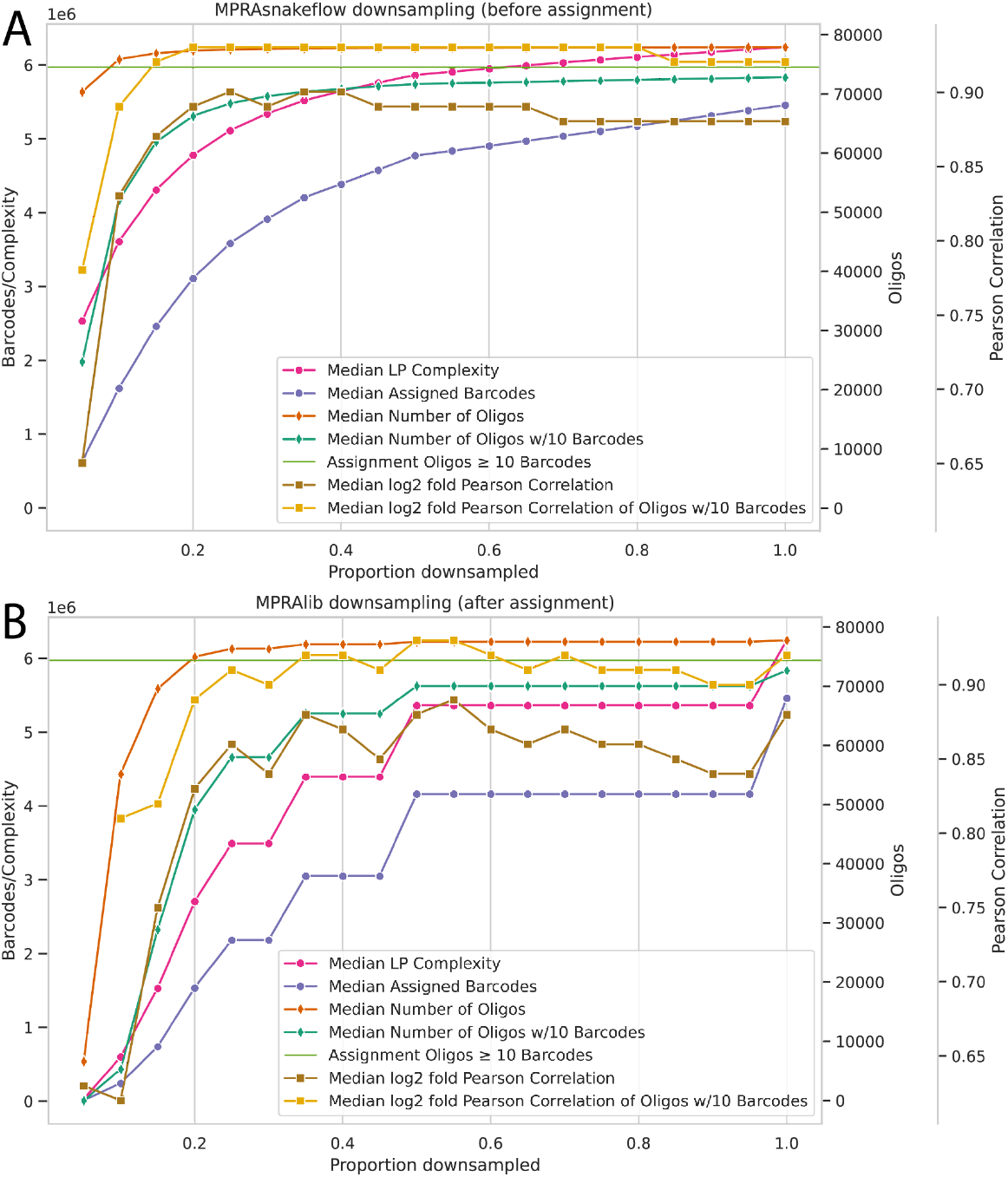
Downsampling analysis for 80K-neurons dataset. DNA and RNA is downsampled in equal proportions to the overall number of counts. Median complexity (Lincoln index), observed assigned barcodes as well as retained oligos (all or with at least 10 barcodes) and Pearson correlation of log2 fold activity (all or oligos with at least 10 barcodes). The horizontal green line shows the number of oligos with at least 10 barcodes in the assignment. (A) RNA and DNA count downsampling in MPRAsnakeflow. This is done before the assignment on the raw counts. (B) Downsampling using the MPRAlib on the overall output (after the assignment).

### Parallel processing and performance benchmarking

We used MPRAsnakeflow (v0.4.4) to analyze a very large experiment (240K-HepG2, see Supplementary Table S1) consisting of ∼240K oligos testing for element-level activity (60K oligos) and variant-level activity (170K oligos). Due to the size of the sequencing files, processing was performed on a high performance computing cluster. Individual job execution was managed using the slurm executor plugin for Snakemake: https://snakemake.github.io/snakemake-plugin-catalog/plugins/executor/slurm.html. Supplementary Table S2 contains a detailed list of MPRAsnakeflow rules with number of jobs, maximum, minimum and average run-time as well as memory usage on the 240K-HepG2 dataset. In summary for the assignment workflow, the three primary input fastq files (paired oligo reads and one barcode read) were split into 30 partitions (`assignment_fastq_split`) resulting in an association pipeline workflow consisting of 164 jobs, with four steps accounting for 150 of these jobs: `assignment_attach_idx`, 60 jobs; `assignment_mapping_bbmap`, 30 jobs; `assignment_mapping_bbmap_getBCs`, 30 jobs; `assignment_merge`, 30 jobs. With a maximum number of parallel jobs set to 24, the workflow completed in under 6 hours with a maximum of 28 GB RAM required for a single job. The majority of jobs for the assignment step required less than 10 GB of RAM and completed in under an hour.

The experiment workflow consisted of 103 jobs and required significantly more time due to only six jobs, specifically the `counts_umi_create_BAM` jobs for the DNA- and RNA-seq files (three of each). The DNA-seq fastq files took ∼12 hours to process while the larger RNA-seq fastq files required ∼34 hours, however these were all run in parallel. The remaining jobs required only four additional hours in total.

### Barcode discrepancy detection and outlier removal

We applied three outlier detection methods to identify problematic barcodes across eight MPRA datasets, comprising six lentiviral-based assays and two episomal-based assays (indicated by “_P” suffix). The three outlier detection methods included: (1) global outliers based on z-score thresholds across all barcodes, (2) oligo-specific outliers based on deviation from within-oligo expression patterns, and (3) expression outliers with extreme log2 expression ratios. Outlier detection patterns revealed noticeable differences between lentiviral and plasmid-based MPRA platforms (Figure 4A). Global outlier detection showed platform-specific bias, with lentiviral assays exhibiting substantially higher rates (0.67-1.56%) compared to plasmid-based assays (0.12-0.48%). Oligo-specific outliers are quite consistent across all experiments, ranging from 0.85% to 1.86%. Notably, large expression outliers are very low in lenti-based assays (up to 0.01%) but had higher rates in plasmid-based assays, in particular 0.53% for 20K_HepG2_P (0.05% for 60K_A549_P). This represents roughly 5-to 53-fold increases over lentiviral datasets.

**Figure 4:**
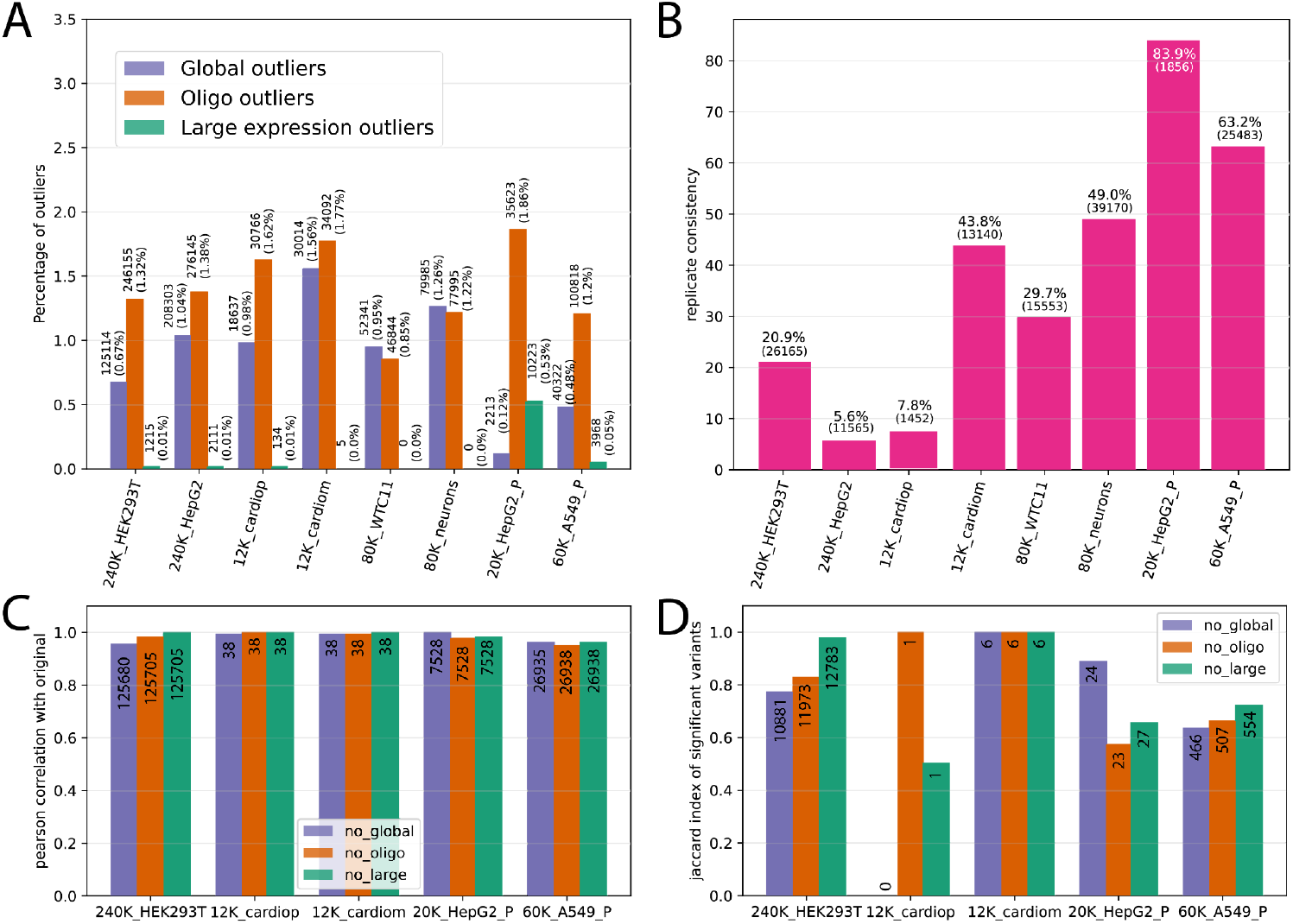
(A) Outlier detection rates on different datasets using three different methods - global outliers (blue), oligo outliers (orange) and expression outliers (green). Percentage shown is the average among the replicates for global and oligo outliers. (B) Consistency of the global outlier detection method across replicates. All lenti-based assays have three replicates, and the 20k and 60k episomal libraries have 6 and 5 replicates respectively. (C) Pearson correlation of variant coefficients between log-fold change estimates from original datasets versus datasets with outliers removed using three methods: global outlier removal (no_global), oligo-specific outlier removal (no_oligo), and large expression outlier removal (no_large). Number of variants per comparison are within the bars. (D) Jaccard indices measuring overlap of significant variants detected in original versus outlier-filtered datasets. Number of overlapping significant variants are written within the bars.

We also detected a difference in the barcode activity distribution (log2 ratio) between lenti and episomal assays. Global outliers for episomal showed a high activity, similar to large expression outliers (see Supplementary Figure S6A).

To assess the reproducibility of MPRA measurements, we examined replicate consistency by calculating the percentage of barcodes that were consistently identified as outliers across all three biological replicates using the global outlier detection method. Replicate consistency varied dramatically across datasets, ranging from 5.6% to 83.9% (Figure 4B). While most lentiviral assays demonstrated poor to moderate consistency (5.6-49.0%), plasmid-based assays exhibited substantially higher consistency rates (63.2-83.9%). This higher consistency in plasmid assays may reflect more uniform transfection conditions and reduced technical noise from integration-site effects that can vary between lentiviral replicates. We also notice an increased replicate consistency observed in differentiated cell types (cardiomyocytes vs. cardioprogenitors: 43.8% vs. 7.8%; neurons vs. WTC11: 49.6% vs. 29.7%) despite expected greater cellular heterogeneity in the former from incomplete differentiation. This may reflect more synchronized transcriptional states and reduced stochastic gene expression noise after differentiation. Additionally, differentiated cells may exhibit more stable chromatin landscapes and consistent regulatory network activity compared to progenitor cells, which are inherently more plastic and variable in their transcriptional responses.

To investigate potential sequence-based biases in outlier detection, we analyzed the GC content distribution of barcodes identified as large expression outliers compared to background barcodes (those with no outliers detected by any method) (Supplementary Figure S6B). Lentiviral-based assays (240K-HEK293T, 240K-HepG2) showed no significant difference in GC content between large expression outliers and background barcodes, with both populations displaying similar median GC content and overlapping distributions. In contrast, plasmid-based assays exhibited divergent patterns. The 20K-HepG2_P dataset showed a significant elevation in GC content among large expression outliers compared to background controls. These results suggest that GC content bias in large expression outliers is not universal but potentially depends on specific experimental conditions, library characteristics or cell-type-specific interactions with GC-rich sequences. The absence of GC bias in lentiviral assays indicates that integration-based delivery may be less susceptible to sequence composition effects compared to episomal approaches.

To assess whether outlier detection is driven by specific problematic sequences or random technical noise, we analyzed barcode sequence overlap between datasets and the concordance of outlier calls for shared sequences (see Supplementary Figure S7). The 15-nucleotide barcode space used in most datasets is enormous (4^15 ≈ 1 billion possible sequences), resulting in minimal sequence overlap between independent libraries. The datasets derived from the same underlying libraries showed substantial overlap: 240K-HEK293T vs 240K-HepG2 (16.2 million shared sequences), 12K-cardiop vs 12K-cardiom (1.9 million shared), and 80K-WTC11 vs 80K-neurons (5.4 million shared). Despite this substantial sequence overlap in related datasets, outlier concordance was remarkably low. For example, for 240K-HEK293T and 240K-HepG2, only 3 out of 16.2 million sequences were identified as large expression outliers in both datasets (Supplementary Figure S7B). This discordance between sequence overlap and outlier concordance indicates that outliers are predominantly driven by dataset-specific technical factors rather than inherent sequence properties.

To assess whether outlier removal improves the quality of MPRA variant effect analysis, we compared results from BCalm for original datasets and datasets with outliers removed. We evaluated two key metrics across five datasets: Pearson correlation of log-fold changes and Jaccard indices measuring the overlap of significant variants detected by each approach. The correlation analysis (Figure 4C) showed consistently high correlations (>0.95) across datasets for all three outlier removal methods, indicating that outlier removal had minimal impact on overall effect size estimates. Similarly, the Jaccard index analysis (Figure 4D) revealed high overlaps in significant variant calls between original and filtered datasets. Eight out of fifteen comparisons showed Jaccard indices above 0.75, with five datasets achieving near-perfect overlap (>0.95).

## Discussion/Conclusion

In this study, we present a comprehensive suite of tools and standards for the uniform processing, analysis, and sharing of MPRA data within the IGVF Consortium and for the MPRA scientific community. Our main contributions are the establishment of harmonized file formats and community standards to facilitate data integration and interoperability, the development of the MPRAsnakeflow workflow for robust, scalable, and reproducible MPRA data processing and the MPRAlib Python library for interactive downstream analysis. Making use of such an unified pipeline, we showcase the processing of data sets with varying design strategies and from different experimental groups. The adoption of standardized file formats for design, counts, effects, and genomic mapping, together with schema validation tools in MPRAlib, represents a major advance for data sharing, reproducibility, and meta-analysis in the field. These standards enable seamless integration of data across experiments, labs, and consortia, and lay the foundation for large-scale comparative studies and benchmarking of regulatory variant effects.

The benchmarking of mapping strategies across diverse MPRA libraries underscores the critical impact of mapping approaches on downstream analyses, particularly for variant-rich designs with small edit distances between sequences. Naive mapping approaches can lead to substantial loss of variant assignments, reducing the quantifiable effects and the interpretability of the assays. We suggest BBmap as a sensible default for MPRAsnakeflow, but enable the easy benchmarking of mapping strategies and suggest it as a best practice in MPRA data analysis.

Our systematic evaluation of barcode outlier detection and removal reveals that while outlier rates and replicate consistency vary significantly across assay platforms and cell types, the impact of outlier removal on downstream variant effect discovery was minimal. We found plasmid-based (episomal) assays to exhibit higher GC-content bias and greater consistency of outlier barcodes across replicates compared to lentiviral assays, which are more susceptible to integration site effects and technical noise. Interestingly, differentiated cell types displayed higher replicate consistency for outlier detection despite greater cellular heterogeneity, suggesting that differentiation may stabilize reporter regulatory activity and reduce technical noise. Based on the overall small effects on activity estimation and differential effects, we suggest that explicit outlier removal is not necessary for variant effect analysis when using BCalm.

While our standardization and implementation in MPRAsnakeflow and associated tools provide a major advance and support a wide range of experimental designs, they are not yet optimized for single-cell or long-read MPRA data, which are emerging areas in regulatory genomics. As a result, we are actively working to extend the support for long-read sequencing and to further optimize the pipeline for new assay modalities. As MPRA designs become more complex and library sizes continue to grow, future work will be needed to reduce technical confounders and adapt to new settings. For example, the field will benefit from a comprehensive database of validated positive controls in multiple tissues and shared reference libraries to enable cross-lab benchmarking and longitudinal tracking of assay performance.

In conclusion, the tools and practices presented here provide a robust, reproducible, and scalable framework for MPRA data analysis, facilitating the functional interpretation of non-coding genetic variation. By enabling consistent data processing and integration, these resources empower researchers to uncover the regulatory architecture of the genome and to accelerate the discovery of causal variants underlying human disease and phenotypic diversity. We anticipate that broad adoption of these standards and pipelines will catalyze collaborative efforts and drive new insights in regulatory genomics.

## Data access

Software and workflow pipelines developed as part of this work include MPRASnakeflow (https://github.com/kircherlab/MPRAsnakeflow), MPRAlib (https://github.com/kircherlab/MPRAlib), and MPRAOligoDesign (https://github.com/kircherlab/MPRAOligoDesign). MPRA datasets used for analysis, together with their portal IDs and dataset names, were obtained from the IGVF portal (http://data.igvf.org) and include IGVFDS1589ODOW (240K-HepG2 and 240K-HEK293T), IGVFDS1419ZPHD (80K-neurons and 80K-WTC11), IGVFDS5307RCQG (20K-HepG2_P), IGVFDS8114HMGD (12K-cardiop and 12K-cardiom), and IGVFDS9668QVZD (8K-neurons). Additionally, data from the ENCODE portal (https://www.encodeproject.org) were analyzed, including ENCSR273YGD (60K-A549_P).

## Competing interest statement

The authors declare no other competing financial interests.

## Acknowledgments

This work has been supported by the Impact of Genomic Variation on Function (IGVF)/NHGRI consortium [UM1 HG011966 /UM1 HG012003] and by the Deutsche Forschungsgemeinschaft (DFG) [464313370]. Computation has been performed on the HPC for Research cluster of the Berlin Institute of Health and the OMICS HPC cluster of the University of Lübeck. We thank all members of the IGVF MPRA focus group for their valuable discussions. We also thank all data producers, especially Beniamin Krupkin, Chengyu Deng, Jessica D. West, Nadav Ahituv, Nicholas F. Page, and Won Ma.

## Author contributions

M.S. and M.L. conceived the project. K.S., J.R., and M.S. performed the mapping benchmarks. J.R. performed the performance benchmarks. N.L. and M.S. performed the complexity analysis. A.V. performed the outlier detection analysis. M.S. developed MPRAlib.

All authors developed or contributed to MPRAsnakeflow and standardized file formats. M.S., M.L., J.R., A.V., K.S., N.L. and M.K. wrote the manuscript.

## Notes

### Competing Interest Statement

The authors have declared no competing interest.

### Summary of Updates

Because the manuscript was directly transferred, no supplemental material is available. The new version consists of the main manuscript along with the uploaded supplemental materials.

