## Supplemental Notes, Figures and Tables for "Uniform processing and analysis of IGVF massively parallel reporter assay data with MPRAsnakeflow"

### Supplemental Material

#### Supplementary Note S1

Here we specify the standardized file formats around experiments. We group them into four categories. 1. Design, 2. Counts 3. Effects and 4. Genomic effects

##### Design

###### Reporter Sequence Design

**Description:** Description of the MPRA design.
**Type:** Tab separated text with headers and without comments. Gzipped.
**Ending:** `.tsv.gz`
**Format:** custom
**Note:** * indicates required and not allowed to be empty, order of the arrays within one entry have to match (first SPDI matches first variant_pos,...), empty arrays are denoted as NA ([] or [NA] or empty string are not allowed).
**Specification:**

```
table reporter_sequence_design
"MPRA sequence design metadata format"
(
string    name*;          “A unique-within-file identifier. Starts with alphabetical
                          characters.”
string    sequence*;      “DNA string of the designed sequence. Allowed chars:
                          [A,T,G,C]”
enum      category*;      “Category of designed sequence. Allowed enums: [variant,
                          element, synthetic, scrambled]”
enum      class*;         “Class of designed sequence. Allowed enums: [test, variant
                          positive control, variant negative control, element active
                          control, element inactive control]”
string    source;         “Free form (e.g. ‘Cardio FG 2022’)."
string    ref;            “Reference genome or sequence (e.g. ‘GRCh38’).”
string    chr;            “Chromosome or contig.”
unit      start;          “0-based position of the left-most position of sequence wrt
                          the reference.”
uint      end;            “1-based position of the right-most position of sequence
                          wrt the reference.”
enum      strand;         “Strand of sequence in reference. Allowed enums: [+,-]
enum[]    variant_class;”What type of variant(s), if applicable. Allowed enums: [SNV,
                          indel]”
Uint[]    variant_pos     “0-based position of the variant(s) starts in the sequence.
                          For indels use normalized representation from SPDI.
                          Acceptable values range from 0 to length(sequence)-1.”
string[]  SPDI            “0-based, validated SPDI representation of the variant(s).”
```

```
enum[]    allele          “The coding with respect to the reference. Allowed enums:
                          [ref, alt]”
string    info            “Any additional comments.”
)
```

#### Reporter Barcode to Element Mapping

**Description:** Creates the link between tested oligos to associated barcodes. Can be pre-designed or learned by association sequencing.
**Type:** Tab separated text without any headers or comments. Gzipped.
**Ending:** `.tsv.gz`
**Format:** custom
**Specification:**

```
table mpra_assignment
"MPRA barcode mapping format"
(
string    barcode;         "Barcode sequence. Allowed chars [A,T,G,C]"
string    oligoName;       "Name of the oligo barcode is assigned to."
)
```

**Example:**

```
GATTTAAACTCGAAT     ID1
CGCGCATCACAACTA     ID2
ATAGCGTAATGGGCA     ID3
TGCTGTTTATGTATG     ID4
AGACTATTTACGAAA     ID5
```

### Counts

#### Reporter Experiment Barcode

**Description:** This format is needed to save the complete measurement on a barcode level of an experiment.
**Type:** Tab separated text with headers and without comments. Gzipped.
**Ending:** `.tsv.gz`
**Format:** custom
**Note:** Columns `dna_count_*` and `rna_count_*` contains a placeholder * for the name of a replicate. E.g. when 3 replicates are used (and names are `1 2 3`) the file format looks like `barcode oligo_name dna_count_1 rna_count_1 dna_count_2 rna_count_2 dna_count_3 rna_count_3`.
**Specification:**

```
table reporter_experiment_barcode
"MPRA experiment barcode level output format"
(
string    barcode;         "barcode, allowed chars [A,T,G,C]"
```

```
string    oligo_name;      "name of the oligo of the design"
uint      dna_count_*      "Number of dna counts of the barcode of a replicate ‘*’, *
                           denotes the name of the replicate (e.g. 1), when no counts
                           observed the entry is empty (or zero)"
uint    rna_count_*        "Number of rna counts of the barcode of replicate ‘*’, *
                           denotes the name of the replicate (e.g. 1), when no counts
                           observed the entry is empty (or zero)"
)
```

**Example:**

```
barcode    oligo_name      dna_count_1  rna_count_1  dna_count_2  rna_count_2
           dna_count_3     rna_count_3
GATTTAAACTCGAAT     ID1                 2      6      1      4
CGCGCATCACAACTA     ID2                 1      2
ATAGCGTAATGGGCA     ID3    12     2
TGCTGTTTATGTATG     ID4                 1      10     1      13
AGACTATTTACGAAA     ID5    2      4     3      1      4      4
```

#### Reporter Experiment

**Description:** This format is needed to save the complete measurement of an experiment.
**Type:** Tab separated text with headers and without comments. Gzipped.
**Ending:** `.tsv.gz`
**Format:** custom
**Specification:**

```
table mpra_experiment
"MPRA experiment output format"
(
string    replicate;       "Name of the replicate."
string    oligo_name;      "Name of the oligo of the design."
uint      dna_counts       "Number of raw DNA counts."
uint      rna_counts       "Number of raw RNA counts."
float     dna_normalized   "Number of normalized/scaled DNA counts (CPM), 4 decimals."
float     rna_normalized   "Number of normalized/scaled RNA counts (CPM), 4 decimals."
float     log2FoldChange   "Fold change (normalized rna/dna ratio, in log2 space),  4
                           decimals."
uint      n_bc             “Number of observed barcodes for the oligo.”
)
```

**Example:**

```
replicate    oligo_name    dna_counts    rna_counts    dna_normalized
rna_normalized      log2FoldChange      n_bc
1      ID3    12     2      8.5714 3.3333 2.5714 1
1      ID5    2      4      1.4285 6,6666 0.2142 1
2      ID1    2      6      2.8571 3.1578 1.1052 1
2      ID2    1      2      1.4285 1.0526 0.7368 1
2      ID4    1      10     1.4285 5.2631 3.6843 1
2      ID5    3      1      4.2857 5.2631 1.2280 1
3      ID1    1      4      1.6666 1.9047 1.1428 1
3      ID4    1      13     1.6666 6.1904 3.7143 1
3      ID5    4      4      6.6666 1.9047 4.0001 1
```

### Effects

#### Reporter Element

**Description:** This format stores the raw format statistical activity analysis for elements. It is dependent on a background/negative set distribution.
**Type:** Tab separated text with headers and without comments. Gzipped.
**Ending:** `.tsv.gz`
**Format:** custom
**Specification:**

```
(
string   oligo_name;        "Name of tested oligo"
float    log2FoldChange;    "Fold change (normalized output/input ratio, in log2
                            space)"
float    inputCount;        “Input count (DNA), normalized (CPM), mean across
                            replicates”
float    outputCount;       “Output count (RNA), normalized (CPM), mean across
                            replicates”
float    minusLog10PValue; "-log10 of P-value"
float    minusLog10QValue; "-log10 of Q-value (FDR)"
)
```

#### Reporter Variant

**Description:** This format stores the raw format statistical activity analysis for variants.
**Type:** Tab separated text with headers and without comments. Gzipped.
**Ending:** `.tsv.gz`
**Format:** custom
**Specification:**

```
(
string   variant_id;        "Variant ID in Canonical SPDI format"
float    log2FoldChange;    "Fold change (alt output/input ratio divided by ref
                            output/input ratio, in log2 space)"
float    inputCountRef;     "Input count reference allele, normalized (CPM), mean
                            across replicates"
float    outputCountRef;    "Output count reference allele, normalized (CPM), mean
                            across replicates"
float    inputCountAlt;     "Input count alternative allele, normalized (CPM), mean
                            across replicates"
float    outputCountAlt;    "Output count alternative allele, normalized (CPM), mean
                            across replicates"
Float    minusLog10PValue; "-log10 of P-value"
float    minusLog10QValue; "-log10 of Q-value (FDR)"
float    postProbEffect;    "Posterior probability of a regulatory effect"
float    CI_lower_95;       "Lower bound of a 95% interval for the variant effect"
float    CI_upper_95;       "Upper bound of a 95% interval for the variant effect"
```

```
uint     variantPos;       “0-based position of the start of the variant in the tested
                           sequence  -1 if aggregation of multiple positions within
                           tested sequences.”
string   refAllele;        “normalized Canonical SPDI reference variant sequence,
                           allowed chars [A,T,G,C,0]. 0 if SPDI reference allele is
                           empty”
string   altAllele;        “normalized Canonical SPDI alternative variant sequence,
                           allowed chars [A,T,G,C,0]. 0 if SPDI alternative allele is
                           empty”
)
```

### Genomic Effects

#### Reporter Genomic Element

**Description:** Defines the activity of an element/region within a genome. Can only be used when exact chromosome start and end location within a reference genome is available. Cannot be used for shuffled or modified elements.
**Type:** Tab separated text without comments and/or headers. Bgzipped
**Ending:** `.bed.gz`
**Format:** BED6+5
**Specification:**

```
table reporter_genomic_element
"BED6+5 MPRA element level common file format"
(
string   chrom;            "Reference sequence chromosome or scaffold"
uint     chromStart;       "Start position in chromosome, 0-based inclusive"
uint     chromEnd;         "End position in chromosome, 0-based exclusive"
string   name;             "Name of tested element or region"
uint     score;            "Indicates how dark the peak will be displayed in the
                           browser (0-1000)"
enum     strand;           "+ or - for strand, . for unknown"
float    log2FoldChange;   "Fold change (normalized output/input ratio, in log2
                           space)"
float    inputCount;       "Input count (DNA), normalized (CPM), mean across
                           replicates"
float    outputCount;      "Output count (RNA), normalized (CPM), mean across
                           replicates"
float    minusLog10PValue; "-log10 of P-value"
float    minusLog10QValue; "-log10 of Q-value (FDR)"
)
```

#### Reporter Genomic Variant

**Description:** Defines the activity of a variant within a genome. Can only be used when exact chromosome start and end location, reference and alternative sequence (Canonical SPDI normalized format) within a reference sequence is available. Cannot be used when no Canonical SPDI is available for the variant.
**Type:** Tab separated text with comments and/or headers. Bgzipped
**Ending:** `.bed.gz`
**Format:** BED6+13
**Specification:**

```
table reporter_genomic_variant
"BED6+10 MPRA variant level common file format"
(
string   chrom;            "Reference sequence chromosome or scaffold"
uint     chromStart;       "Start position of the variant in chromosome, 0-based
                           inclusive"
uint     chromEnd;         "End position of the variant in chromosome, 0-based
                           exclusive"
string   name;             "Name of tested variant"
uint     score;            "Indicates how dark the peak will be displayed in the
                           browser (0-1000)"
enum     strand;           "+ or - for strand the variant was tested in, . for
                           unknown"
float    log2FoldChange;   "Fold change (alt output/input ratio divided by ref
                           output/input ratio, in log2 space)"
float    inputCountRef;    "Input count reference allele, normalized (CPM), mean
                           across replicates"
float    outputCountRef;   "Output count reference allele, normalized (CPM), mean
                           across replicates"
float    inputCountAlt;    "Input count alternative allele, normalized (CPM), mean
                           across replicates"
float    outputCountAlt;   "Output count alternative allele, normalized (CPM), mean
                           across replicates"
float    minusLog10PValue; "-log10 of P-value"
float    minusLog10QValue; "-log10 of Q-value (FDR)"
float    postProbEffect;   "Posterior probability of a regulatory effect"
float    CI_lower_95;      "Lower bound of a 95% interval for the variant effect"
float    CI_upper_95;      "Upper bound of a 95% interval for the variant effect"
uint     variantPos;       “0-based position of the start of the variant in the tested
                           sequence, -1 if aggregated effect over multiple positions”
string   refAllele;        “normalized Canonical SPDI reference variant sequence,
                           allowed chars [A,T,G,C,0]. 0 if SPDI reference allele is
                           empty”
string   altAllele;        “normalized Canonical SPDI alternative variant sequence,
                           allowed chars [A,T,G,C,0]. 0 if SPDI alternative allele is
                           empty”
)
```

### Supplementary Note S2

All MPRA experiments used in this work with short descriptions. For an overview with references to datasets see Supplementary Table S1.

#### 240K-HepG2 and 240K-HEK293T

Very large library testing variants/regions in open chromatin (screen regions) in front of genes (range between 500 and 20K nt before TSS). Open chromatin regions of HepG2, HEK293T MCF-7 and stem cells are used. Variants where selected from an in-silico mutagenesis of all regions, predicted by an DNN trained on open chromatin in multiple cell-lines. High active, low active variants are selected.

#### 80K-WTC11 and 80K-neurons

Variants/Region design of open chromatin regions that are close to disease-causing genes related to neuronal and cardiac phenotypes. Also for all actionable genes (CAVA). All common variants (>5% AF) as well as a subset, prioritized by Enformer, of rare and singleton variants.

## 60K-A549_P

An episomal library tested in A549 cells, which was part of a MPRA set used to engineer and validate synthetic CREs capable of driving gene expression with programmed cell-type specificity (Gosai et al. 2024). This library was released via ENCODE.

#### 20K-HepG2_P

An episomal library tested in HepG2 cells. Lead GWAS variants for lipid-related traits (HDL-cholesterol, LDL-cholesterol, triglycerides, and the liver enzymes ALT, AST, and GGT) were selected from large meta-analyses (Vujkovic et al. 2022; Sakaue et al. 2021; Pazoki et al. 2021; Gao et al. 2021; Kanoni et al. 2022; Graham et al. 2021). Complete signals were built by identifying variants in high linkage disequilibrium (LD; $r^2 \geq 0.8$ in European reference panel).

#### 12K-cardiop and 12K-cardiom

Small library with mostly regions tested in cardiomyocytes and cardioprogenitor cells derived from WTC11.

#### 8K-neurons

Small library on ASD risk variants. See the original publication for more details (Koesterich et al. 2023).

### Supplementary Tables

**Supplementary Table S1:** Overview of the used MPRA datasets within this work. ID - The used identifier within this manuscript. Assay - The assay type (lentiMPRA or episomal MPRA). Tissue - The name of the conducted tissue within the experiment with the in vitro system ID from the IGVF portal (data.igvf.org). Portal ID - The construct library ID for IGVF data samples (data.igv.org) or the ENCODE ID of the functional characterization experiment (encodeproject.org). #Oligos - Number of design oligos. #Regions - Number of designed regions (unique mappable region within the human reference genome). #Variants - Number of designed variants. Complexity - Shows the estimated barcode complexity by the median Lincoln index across all replicates. PMID - Pubmed identifier of the original publication of the dataset, if available. For 60K-A549_P variants and oligos are estimated from the design file ENCFF074MMO. Supplemental Figures

| ID | Assay | Tissue | Portal ID | #Oligos | #Regions | #Variants | Complexity | PMID |
|---|---|---|---|---|---|---|---|---|
| 240K-HepG2 | lenti | HepG2 IGVFSM9009DVDG | IGVFDS1589ODOW | 238,455 | 64,058 | 170,048 | 20,410,541 | |
| 240K-HEK293T | lenti | HEK293T IGVFSM8026MGEJ | IGVFDS1589ODOW | 238,455 | 64,058 | 170,048 | 19,451,971 | |
| 80K-neurons | lenti | WTC11 ngn2 derived neurons IGVFSM5401ZHCC | IGVFDS1419ZPHD | 80,215 | 30,054 | 46,467 | 6,243,618 | |
| 80K-WTC11 | lenti | WTC11 IGVFSM9779NDRC | IGVFDS1419ZPHD | 80,215 | 30,054 | 46,467 | 5,632,412 | |
| 60K-A549_P | episomal | A549 | ENCSR273YGD | 59,657 | 27,479 | 27,284 | 7,580,455 | 39443793 |
| 20K-HepG2_P | episomal | HepG2 IGVFSM9800GLLF | IGVFDS5307RCQG | 25,026 | 13,397 | 11,959 | 1,588,034 | |
| 12K-cardiop | lent | WTC11 derived cardioprogenitors IGVFSM9821BABW | IGVFDS8114HMGD | 11,928 | 11,511 | 83 | 1,874,896 | |
| 12K-cardiom | lenti | WTC11 derived cardiomyocytes IGVFSM9501XSAN | IGVFDS8114HMGD | 11,928 | 11,511 | 83 | 1,907,408 | |
| 8K-neurons | lenti | neural progenitor cells (Neural induction of H1 hESCs) IGVFSM0720XHJA | IGVFDS9668QVZD | 7502 | 3899/3900 | 3456/3600 | 1,523,572 | 36834916 |

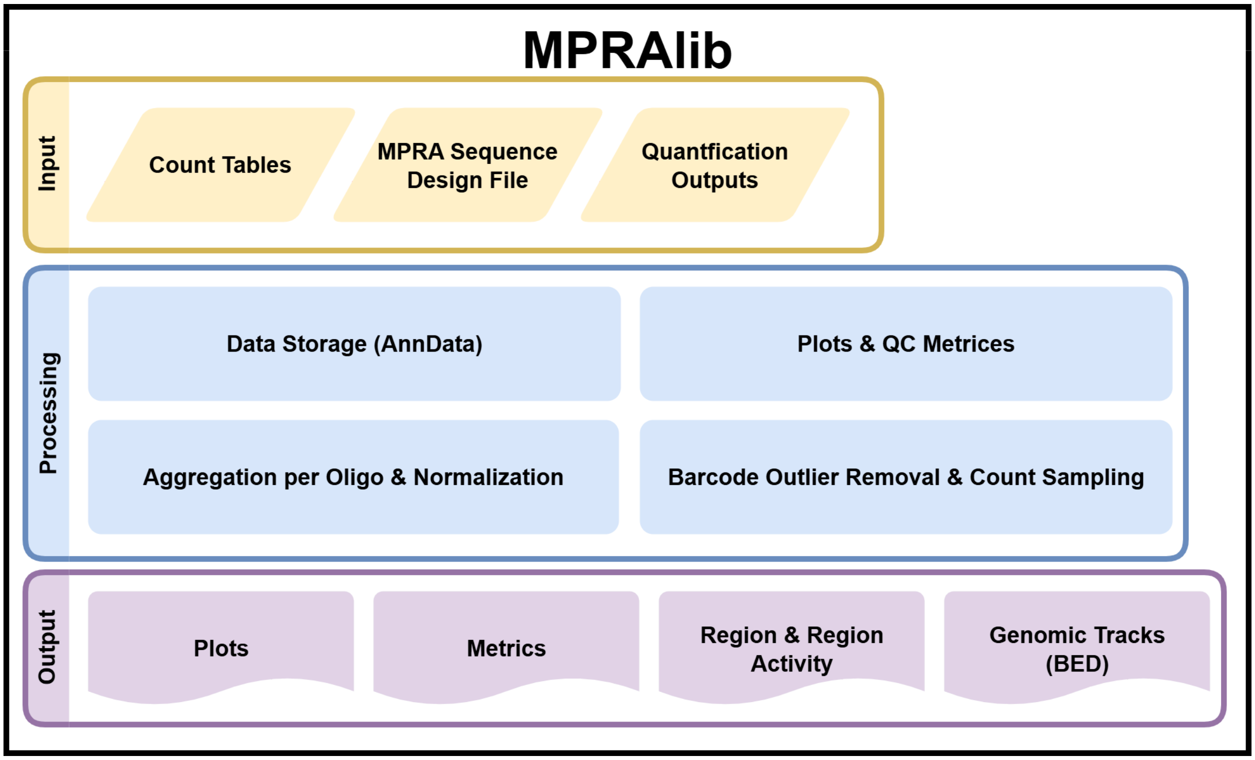


**Supplementary Figure S1:** Overview of the python library MPRAlib. Inputs are shown in yellow, main processing steps in blue, and outputs in purple.

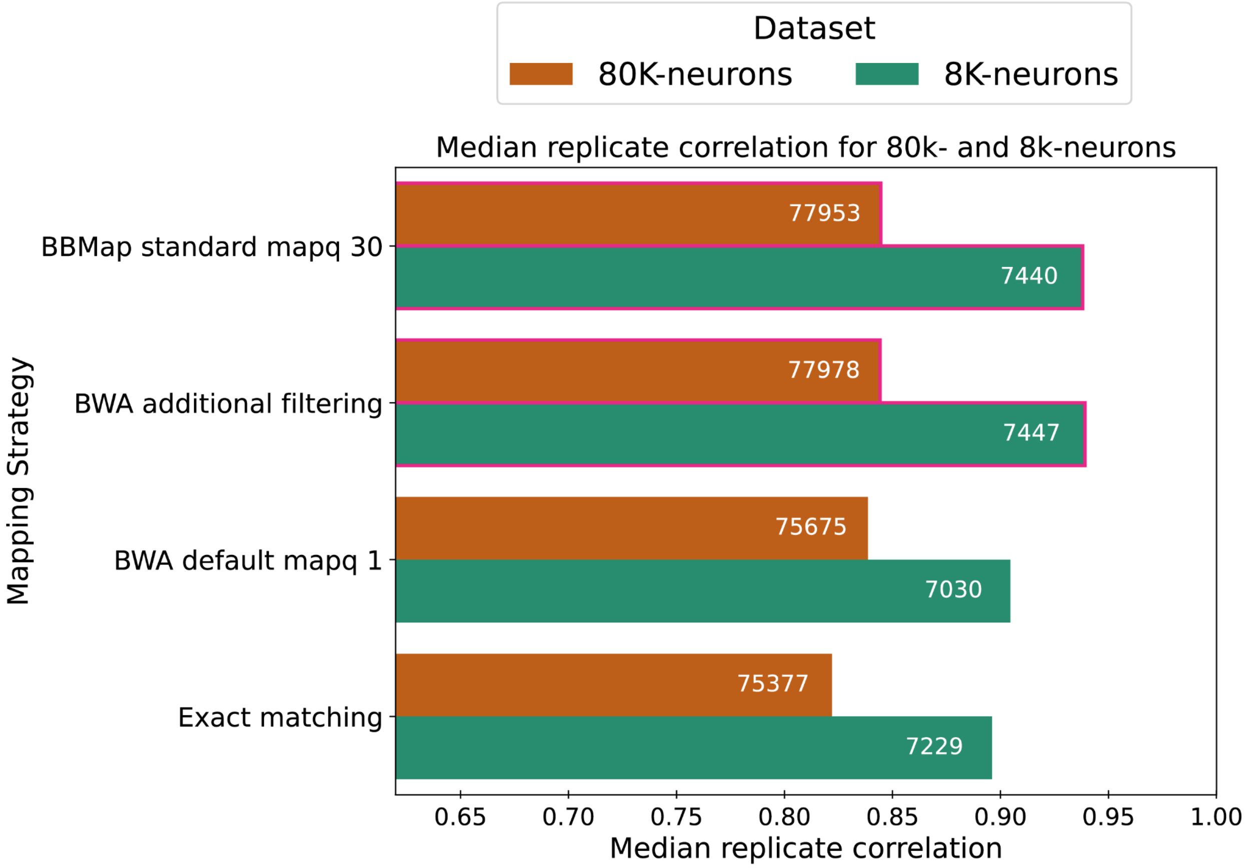


**Supplementary Figure S2:** Median Spearman replicate (n=3) correlation of the oligo activity (log2(RNA/DNA)) for the 8K-neurons (green) and 80K-neurons (orange) dataset. The highlighted bar represents the mapping strategy with the highest correlation (rounded to the second digit).

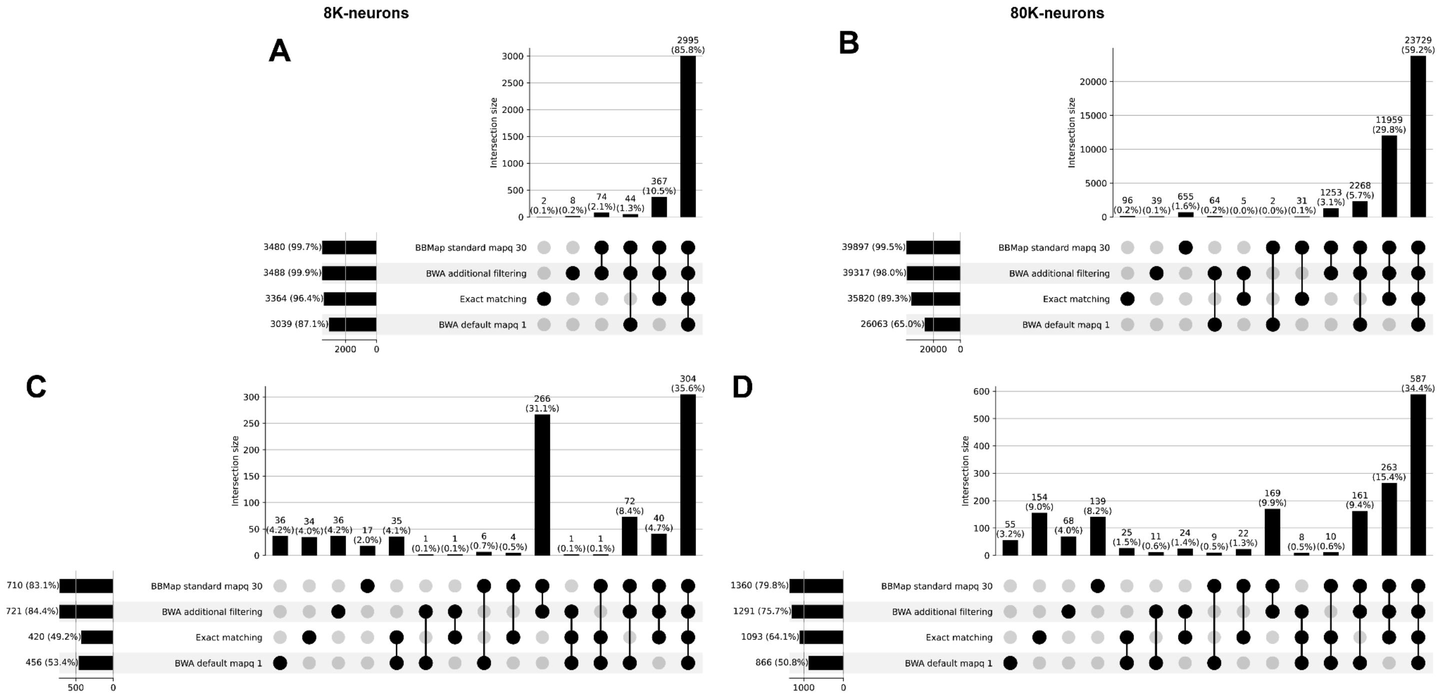


**Supplementary Figure S3:** Comparing BCalm results using 8K-neurons (left panels) and 80K-neurons (right panels) on the four different mapping strategies. Each of the panels contains an upset plot, which shows the set size for each mapping strategy and the size of the respective set overlap on the right. The size of the overlap is represented by the number annotated at the top of the bar. The sets used to obtain the overlap represented by the bar are shown by a black dot in the column below the bar. (C) and (E) represent the results for 8K-neurons with around 3,600 variants, and (D) and (E) show the results of 80K-neurons with around 46,000 variants. (C) and (D) show the variants with a readout between the mapping strategies, while (E) and (F) focus on the overlap between significant variants.

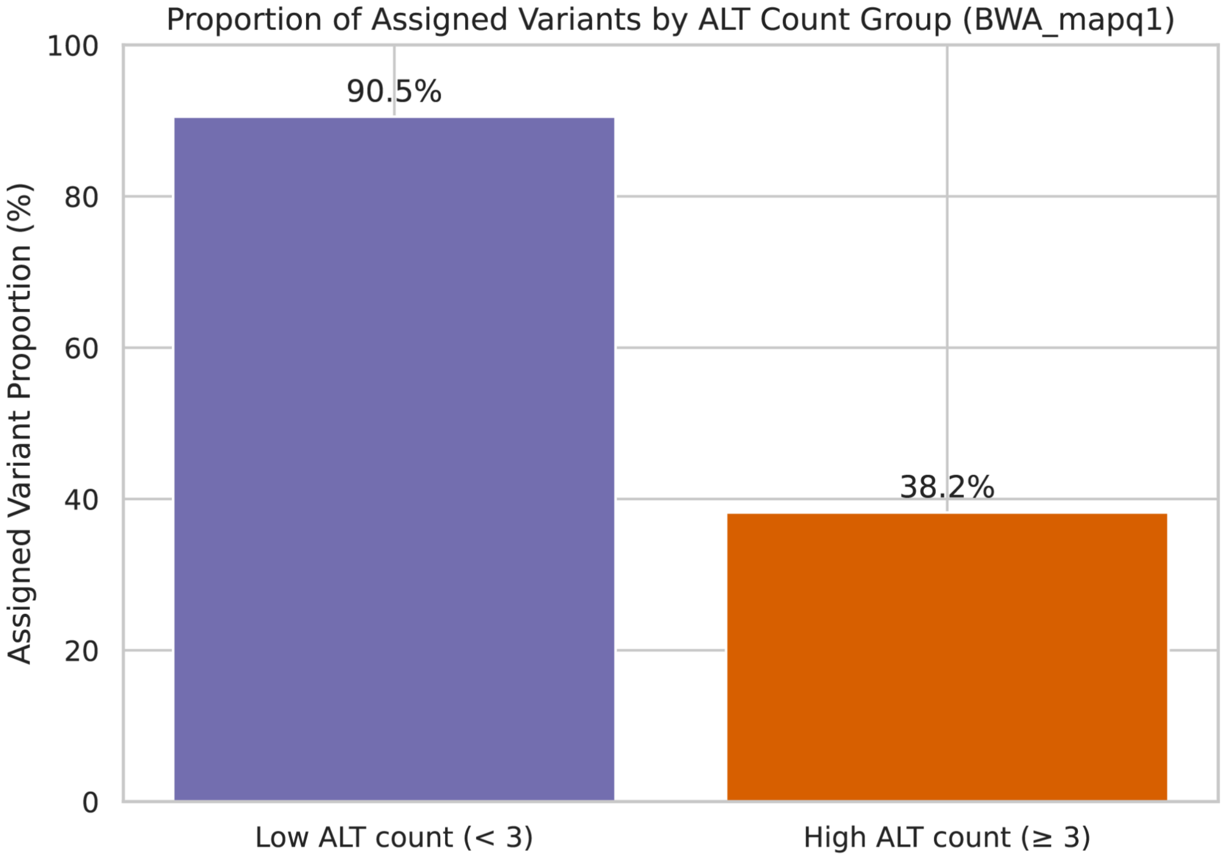


**Supplementary Figure S4:** Number of assigned variants for the 80K-neurons/WTC11 library by MPRAsnakeflow using the BWA default approach. Variants are grouped by the number of ALT alleles that are designed on the same reference sequence. The left bar shows all assigned variants with reference alleles having one or two alternative alleles (ALT) designed (n: 15927/17599). Right side with three or more per reference (n: 10992/28775).

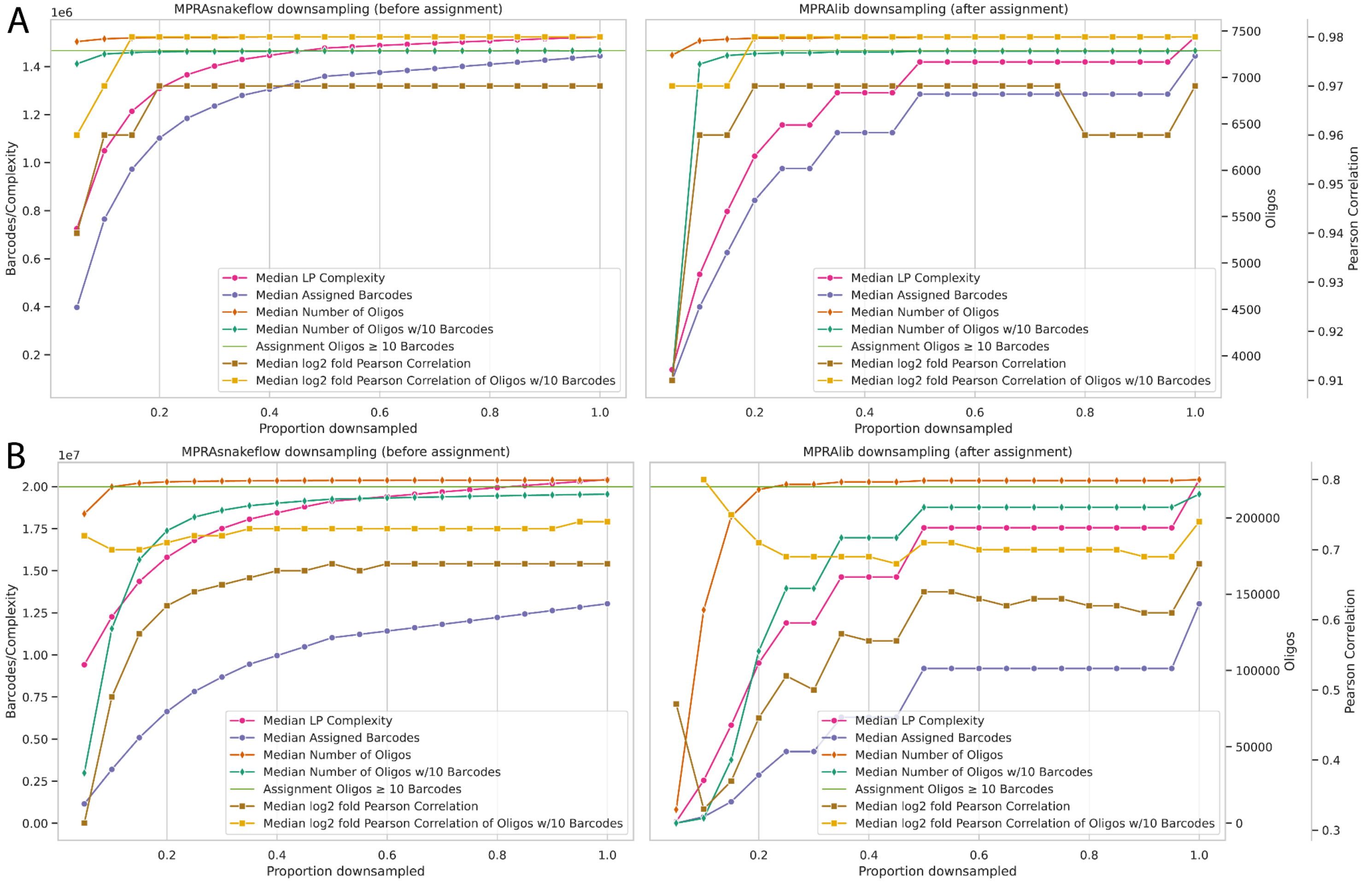


**Supplementary Figure S5:** Downsampling analysis for (A) 8K-neurons, and (B) 240K-HepG. Median complexity (Lincoln index), observed assigned barcodes as well as retained oligos (all or with at least 10 barcodes) and Pearson correlation of log2 fold activity (all or oligos with at least 10 barcodes). The horizontal green line shows the number of oligos with at least 10 barcodes in the assignment. The left side uses RNA and DNA count downsampling in MPRAsnakeflow, which happens before the assignment (raw counts). The right side shows downsampling using the MPRAlib (DNA and RNA to equal proportions).

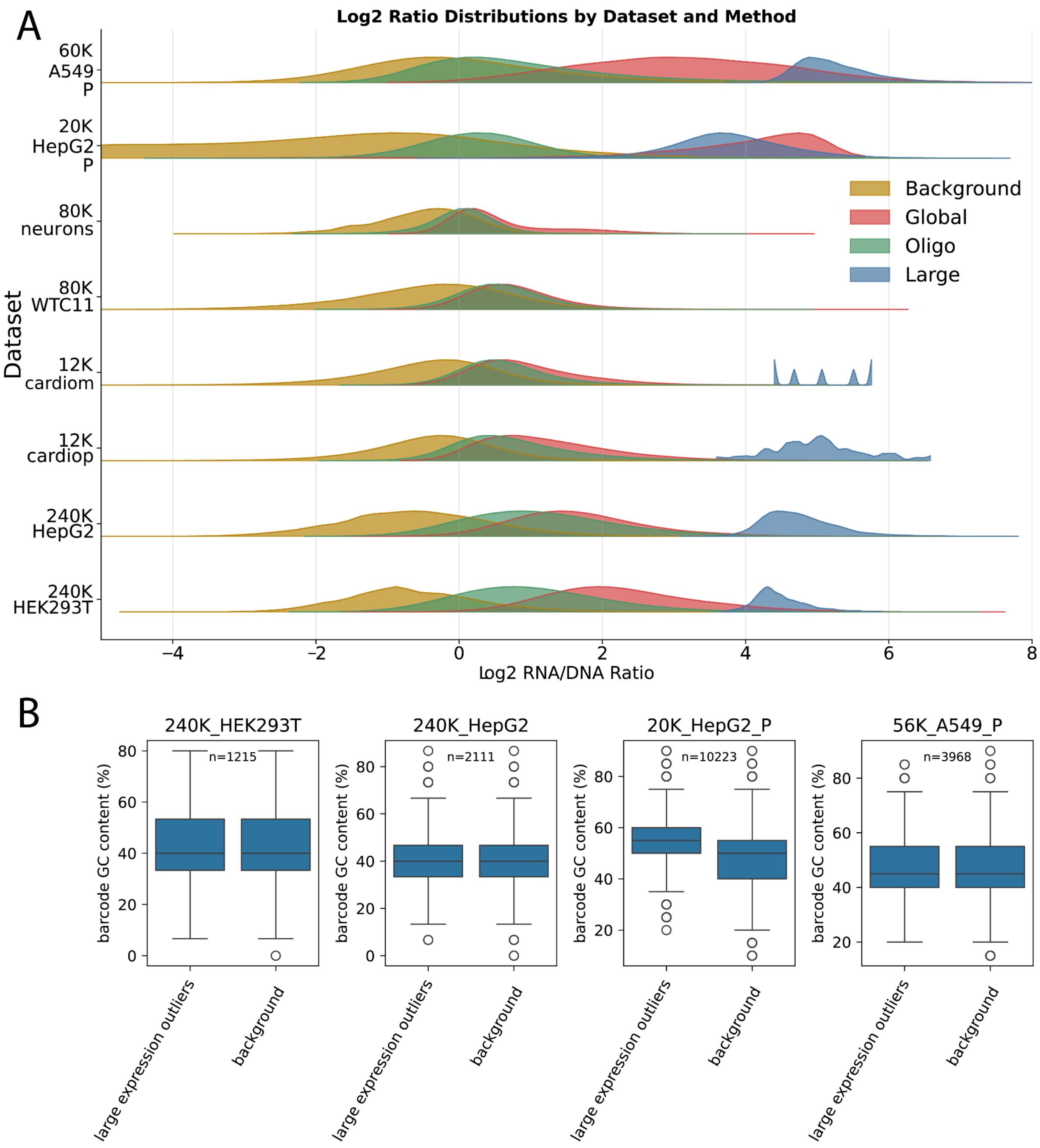


**Supplementary Figure S6:** (A) Distribution of log2 ratios for the eight datasets used for outlier detection grouped as background (all barcodes not flagged by any outlier method, yellow) and different outlier detection methods (global z-score, oligo-specific, and large expression in pink, green, and blue, respectively). (B) Barcode GC content comparison between large expression outliers and the background (all detected barcodes within the library).

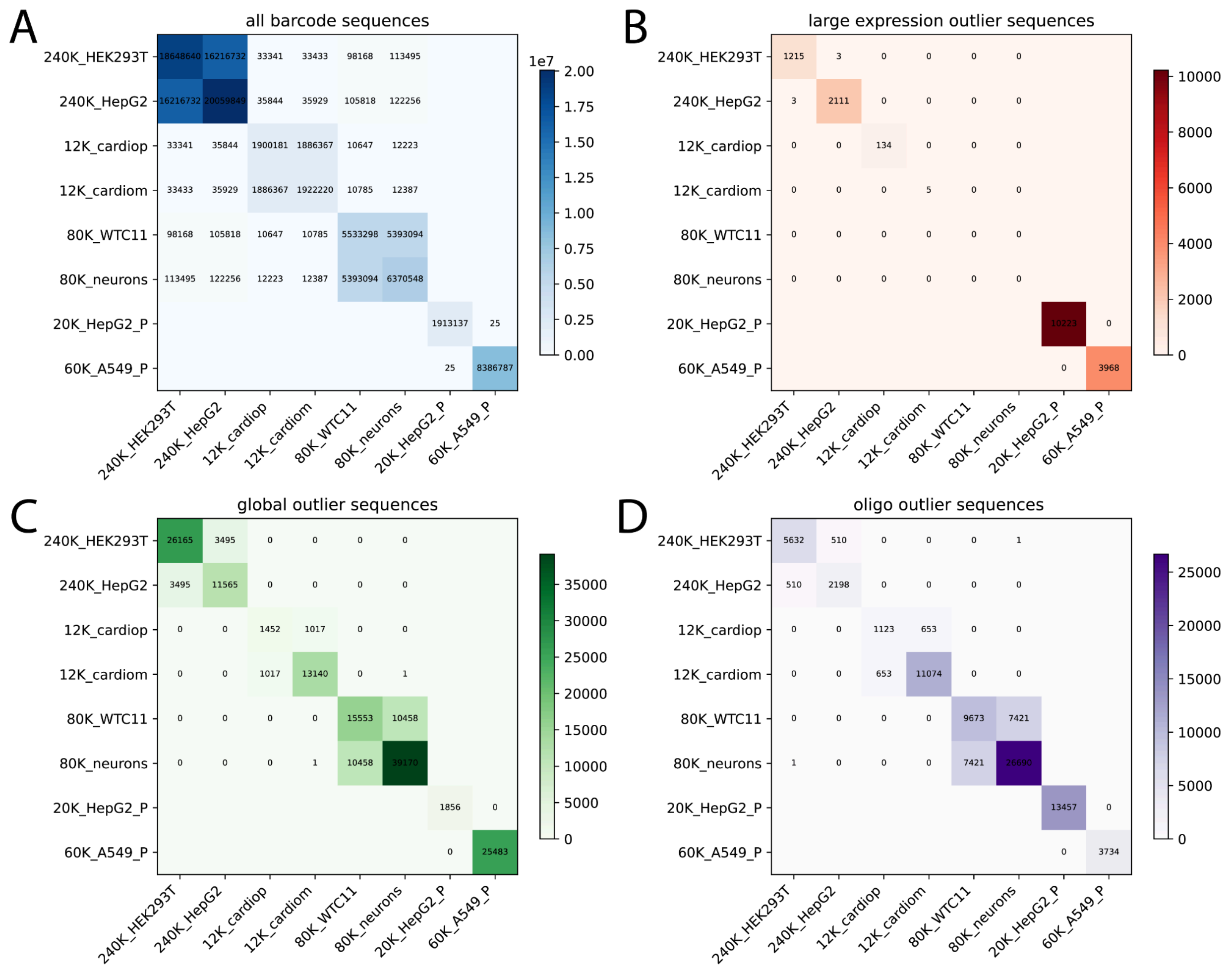


**Supplementary Figure S7:** Intersection matrix for (A) total barcodes detected and outlier barcodes (B - large expression, C - global outliers, D - oligo outliers) across datasets (lenti based libraries have 15 nt barcodes and episomal libraries have 20 nt barcodes).
